# Cognitive control of orofacial and vocal responses in the human frontal cortex

**DOI:** 10.1101/698613

**Authors:** Kep Kee Loh, Emmanuel Procyk, Remi Neveu, Franck Lamberton, William Hopkins, Michael Petrides, Céline Amiez

## Abstract

The frontal cortical areas critical for human speech production, i.e. the ventrolateral frontal cortex (cytoarchitectonic areas 44 and 45; VLF) and the dorsomedial frontal cortex (DMF) comprising the mid-cingulate cortex (MCC) and the pre-supplementary motor area (preSMA), exist in non-human primates and are implicated in cognitive vocal control functions. The present functional neuroimaging study seeks to define the basic roles of these VLF-DMF network regions in primate vocal production and how they might have been adapted for human speech. We demonstrate that area 44 and the MCC are respectively involved in the cognitive selection of orofacial, non-speech vocal and verbal responses, and the feedback-driven adaptation of these responses – roles that are likely preserved across primates. In contrast, area 45 and preSMA have roles that are specific to human speech: area 45 contributes to active verbal retrieval during learning, while preSMA is involved in processing verbal feedback during orofacial/vocal adaptations.

## Introduction

The capacity for speech is uniquely human. Despite centuries of research, the question of how human speech and its neural correlates emerged during primate evolution has remained highly controversial. This is largely attributed to the difficulty in accepting continuity between human speech, which involves the flexible generation and usage of complex vocalizations, and non-human primate (NHP) vocalizations, which appear to be limited to a set of fixed calls that are tied to specific emotional and motivational situations ^1^. However, recent evidence has suggested that volitional and flexible vocal control is indeed present in NHPs ^2^. Importantly, the complexity of cognitive vocal control appears to increase across the primate phylogeny: while monkeys can flexibly initiate calls and switch between them ^3–5^, chimpanzees and orangutans exhibit a higher level of vocal flexibility in generating novel vocalizations and using them in a goal-directed manner ^6, 7^. These findings suggest that human speech, and its associated neural systems, could have evolved from a basic cognitive vocal control system that already exists in NHPs. The identification of this early cognitive vocal control system and its generic functions across primates thus provides a critical step in understanding the emergence of human speech during primate evolution.

Across primates, two anatomically homologous frontal systems are implicated in the cognitive control of vocalizations ^reviewed in 2^: the ventrolateral frontal cortex (VLF), that comprises cytoarchitectonic areas 44 and 45; and the dorsomedial frontal cortex (DMF), that comprises the pre-supplementary area (pre-SMA), supplementary motor area (SMA) and the mid-cingulate cortex (MCC). In humans, the VLF houses the Broca’s region (i.e. areas 44 and 45) which yields severe speech impairments when damaged. Electrical stimulation of area 44, which lies immediately anterior to the ventral precentral motor region involved in the control of the orofacial musculature, results in speech arrest ^8^. Functional neuroimaging studies suggest a role of area 45 in controlled verbal memory retrieval ^9^ which is often expressed in verbal fluency ^10,11^. Electrical stimulation in the human DMF results in vocalization in a silent patient, and speech interference or arrest in a speaking patient ^8^. DMF lesions have been associated with long-term reduction in verbal output ^12^. Importantly, Chapados and Petrides ^13^ further noted that within the DMF, lesions must include SMA, pre-SMA and the MCC regions to induce deficits, suggesting the existence of a local DMF network contributing to vocal and speech production. Intriguingly, even though NHPs do not speak, the anatomical homologues of VLF and the DMF regions exist and are implicated in aspects of cognitive vocal control ^reviewed in^ ^2,14^ In monkeys, VLF neurons show increased firing rates prior to voluntary call initiations ^3^ and electrical stimulations in this region (specifically, in area 44) evoke orofacial movements ^15^. In chimpanzees, increased VLF activity is observed during the production of intentional attention-getting vocalizations ^16^. The DMF regions are also crucial for voluntary vocal productions: stimulations in the anterior MCC and around the pre-SMA/SMA border evoke orofacial movements and vocal productions ^17–19^, while lesions in these regions result in reduced spontaneous vocalizations ^20,21^ and abolish the production of conditioned calls ^22,23^. As such, this VLF-DMF brain network could constitute a primitive cognitive vocal control system that developed during primate evolution to support the expanded vocal flexibility present in humans. Currently, however, it is virtually unknown how the various VLF and DMF regions contribute to the cognitive control of vocalizations in primates and how they might have changed across primate evolution.

The present study seeks to disentangle the individual roles of VLF-DMF regions in cognitive vocal control that might be generic across primates. In both humans and macaques, the posterior lateral frontal cortex that lies immediately anterior to the precentral motor zone, where area 44 is found, has been linked to the cognitive selection between competing motor acts ^14^; whereas the MCC is typically associated with behavioral feedback evaluation during learning ^25,28,29^. On this basis, we hypothesize that area 44 is involved in the high-level cognitive selection between competing orofacial and vocal acts, while the MCC is involved in the use of vocal feedback for adapting vocal behaviors. To capture these processes, we utilized a conditional associative learning protocol that requires subjects to first learn, via trial-and-error with vocal feedback, to select particular responses from a pool of available responses based on if – then relations. We examined the brain networks involved in selecting a) orofacial, b) non-speech vocal, c) speech vocal and d) manual responses based on external visual cues, and in processing vocal feedback to drive trial-and-error learning.

Our results provided major novel insights into the contributions of the VLF-DMF network regions in orofacial, nonspeech vocal and verbal production: Area 44 is involved in the cognitive selection of orofacial, nonspeech vocal, and verbal responses (but not manual responses) conditional upon learned relations to external stimuli. In contrast, area 45 is specifically recruited for the selection of nonspeech vocal and verbal responses during learning, i.e. when the if - then conditional relations had not yet been mastered and, therefore, the active cognitive mnemonic retrieval load was high. The MCC is involved in processing nonspeech vocal and verbal feedback, and effector-independent cognitive response selection during learning, and the preSMA is selectively involved in the cognitive control of verbal response selections based on verbal feedback.

## Results

Subjects underwent three fMRI sessions in which they performed the visuo-motor conditional associative learning task (Fig 1A) and control task (Fig 1C) with different motor responses (Fig 1B): Manual (button presses), Orofacial (mouth movements), and Vocal (either nonspeech or verbal vocalizations). In each learning task block (Fig 1A), subjects had to learn the correct conditional pairings between three different motor responses and visual stimuli based on the non-speech vocal or verbal feedback provided (learning phase), and subsequently perform the learnt associations (post-learning phase). In each control task block (Fig 1C), subjects performed an instructed response to three possible visual stimuli.

**Figure 1.**
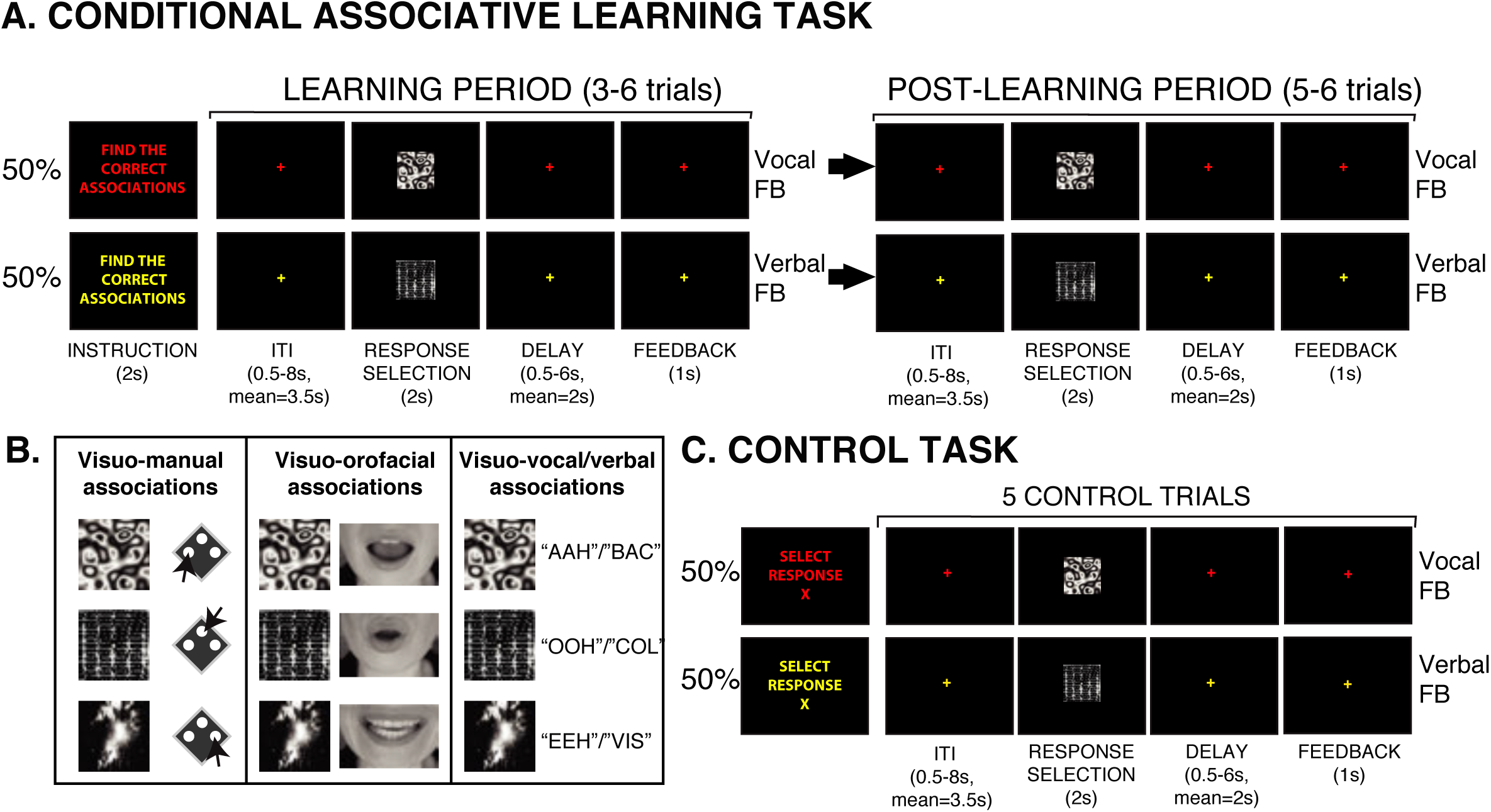
Experimental tasks. A. Conditional associative learning task. In the learning phase, subjects have to discover, by trial-and-error, the correct pairings between three possible motor responses and three visual stimuli. In each trial, one visual stimulus was randomly presented for 2 seconds, in response to which the subjects selected and performed one of the three motor responses (Response Selection). If the instruction font and fixation cross were red (in 50% of learning sets), vocal feedback was provided to indicate whether the response was correct (“AHA”) or wrong (“BOO”). If the instruction font and fixation cross were yellow (in 50% of learning sets), verbal feedback was provided to indicate whether the response was correct (“CORRECT”) or wrong (“ERROR”). After a subject had performed a correct response to each one of the three stimuli (which indicated the end of the learning period), the subjects had to perform each of learnt associations twice (i.e. post-learning period). B. Visuo-motor associations that subjects had to learn in the 3 versions of the visuo-motor conditional associative learning task. In the visuo-manual condition, subjects learn associations between 3 button presses and 3 visual stimuli. In the visuo-orofacial condition, subjects learn associations between 3 orofacial movements and 3 visual stimuli. In the visuo-vocal/verbal condition, subjects learn associations between 3 vocal (“ahh”, “iih”, “ooh”) or 3 verbal (“bac”, “vis”, “col”) responses and 3 visual stimuli. C. Visuo-motor control task with vocal feedback (50% of trials, indicated by red colored fonts of the instructions and fixation crosses) or verbal feedback (50% of trials, indicated by yellow colored fonts of the instructions and fixation crosses). In the control task, the subjects perform the instructed (X) motor response to every presented stimulus during response selection for 5 consecutive trials. All motor responses used in the conditional associative learning task (the 3 manual, 3 orofacial, and 3 vocal/verbal) are instructed to be performed in control trials over the 3 fMRI sessions.

### Functional dissociations in the posterior lateral frontal cortex for the cognitive selection of manual versus orofacial, nonspeech vocal, and verbal responses

The mean BOLD signal during the response selection epoch was compared between post-learning versus control trials and between learning versus control trials involving manual, orofacial, non-speech vocal and speech vocal responses. In line with previous findings^25^, group-level analyses demonstrated increased BOLD activity in the dorsal premotor region (PMd) during the conditional selection of <u>manual</u> responses in both the learning and post-learning periods relative to control (Fig 3A, see Table S1 for activation peak locations and t-values). Single-subject analyses confirmed that individual PMd peaks, during both the learning (observed in 17/18 subjects) and post-learning phases (observed in 18/18 subjects), were consistently located in the dorsal branch of the superior precentral sulcus. Importantly, no significant activation was observed in the ventrolateral part of the posterior frontal region where area 44 lies, during both the learning and the post-learning period of manual selections (Fig 3A, Table S1).

**Figure 3.**
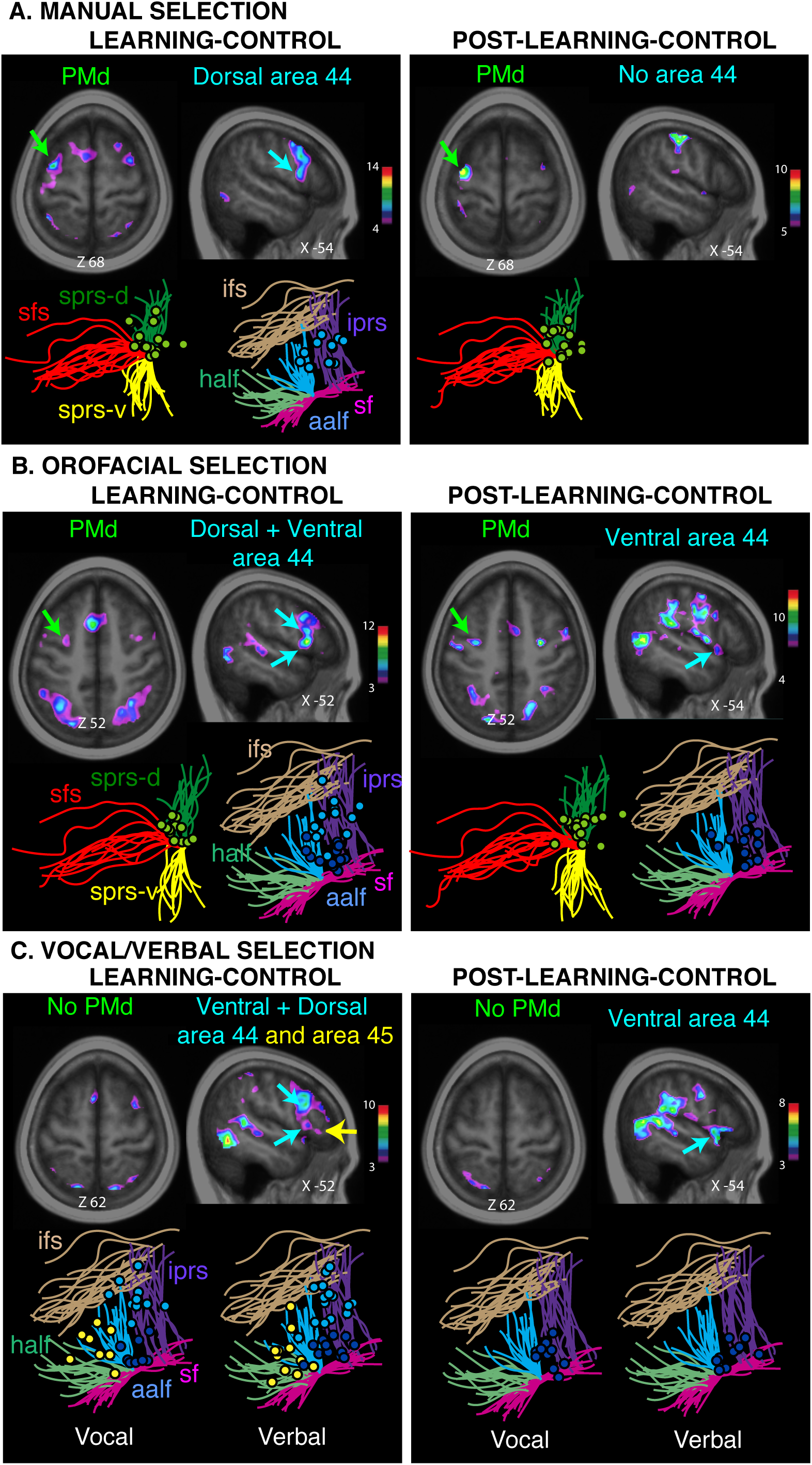
Conditional response selection in the posterior lateral prefrontal cortex. Group (above) and individual subject activations (below) are shown as dots around relevant sulci during (A) manual response selection in the learning and post-learning periods, (B) during orofacial response selection in the learning and post-learning periods, (C) during vocal and verbal response selection in the learning and post-learning periods. Green circles depict individual subject PMd activations for manual and orofacial response selection during learning and post-learning, respectively. Light and dark blue circles depict individual dorsal and ventral area 44 activations and yellow circles depict area 45 activity during response selection. Abbreviations: cs, central sulcus, sprs-d, dorsal superior precentral sulcus, sprs-v, ventral superior precentral sulcus, sfs, superior frontal sulcus, ifs, inferior frontal sulcus, iprs, inferior precentral sulcus, as, ascending sulcus, hs, horizontal sulcus, lf, lateral fissure.

By contrast, orofacial response selection resulted in increased BOLD activity in both ventral area 44 and the PMd during both the learning and post-learning periods relative to control (Fig 3B, see Table S1 for activation peak locations and t-values). Subject-level analyses in the left hemisphere showed that individual ventral area 44 peaks were observed in the pars opercularis (14/18 subjects in both learning and post-learning periods), and the PMd peaks were consistently found in the dorsal branch of the superior precentral sulcus (observed in 14/18 subjects during the learning period and in 17/18 subjects in the post-learning period) (Fig 3B, Table S1).

The selection of vocal responses (pooled across nonspeech vocal and verbal responses in the group analysis) was associated with increased BOLD activity in the left ventral area 44, and not the PMd, during both learning and post-learning, relative to control (Fig 3C; see Table S1 for activation locations and t-values). At the single-subject level, we assessed the brain activations associated with verbal and nonspeech vocal responses separately. We observed that individual ventral area 44 peaks (dark blue circles) during both learning (verbal peaks: 14/18 subjects, nonspeech vocal peaks: 9/18 subjects) and post-learning phases (verbal peaks: 10/18 subjects, nonspeech vocal peaks: 12/18 subjects), were consistently located in the pars opercularis region bounded anteriorly by the ascending ramus of the Sylvian fissure, posteriorly by the inferior precentral sulcus, dorsally by the inferior frontal sulcus, and ventrally by the Sylvian fissure (Fig 3C). This finding indicated that both the verbal and the nonspeech vocal response selections recruted the same ventral area 44.

### Functional dissociations within Broca’s region during cognitive manual, orofacial and vocal selection

Across all response conditions, we observed increased activity in the dorsal part of area 44 as subjects selected their responses during learning (Fig 3A, 3B, 3C; see Table S1 for activation location and t-value), but not during post-learning (Table S1), suggesting that it contributes specifically to the learning of conditional associations, and crucially, in an effector-independent manner. In contrast, ventral area 44 is effector-dependent as it is recruited in the cognitive selection of orofacial, nonspeech and speech vocal responses – but not of manual responses-during both learning and post-learning.

In contrast to the involvement of ventral area 44 in orofacial and both speech and nonspeech vocal conditional response selections, area 45 showed increased activity only for vocal conditional response selections (pooled across nonspeech vocal and verbal responses) during the learning period (see Fig 3C; Table S1). The single-subject level analysis revealed that increased activity in area 45 (yellow circles) during learning (verbal peaks: 11/18 subjects, nonspeech vocal peaks: 9/18 subjects) is consistently located in the pars triangularis region bounded posteriorly by the ascending ramus of the Sylvian fissure, dorsally by the inferior frontal sulcus, and ventrally by the Sylvian fissure (Fig 3C).

### The dorsomedial frontal cortex is involved during the learning of conditional visuo-motor associations

The comparison between the BOLD signal observed at the response selection epochs in learning versus control trials and post-learning versus control trials revealed increased activity in the dorso-medial frontal cortex (DMF) only during the learning, but not post-learning period (Fig 4, Table S2). Thus, contrary to the posterior lateral frontal cortex, the DMF is not involved in the post-learning selection of conditional responses from various effectors.

**Figure 4.**
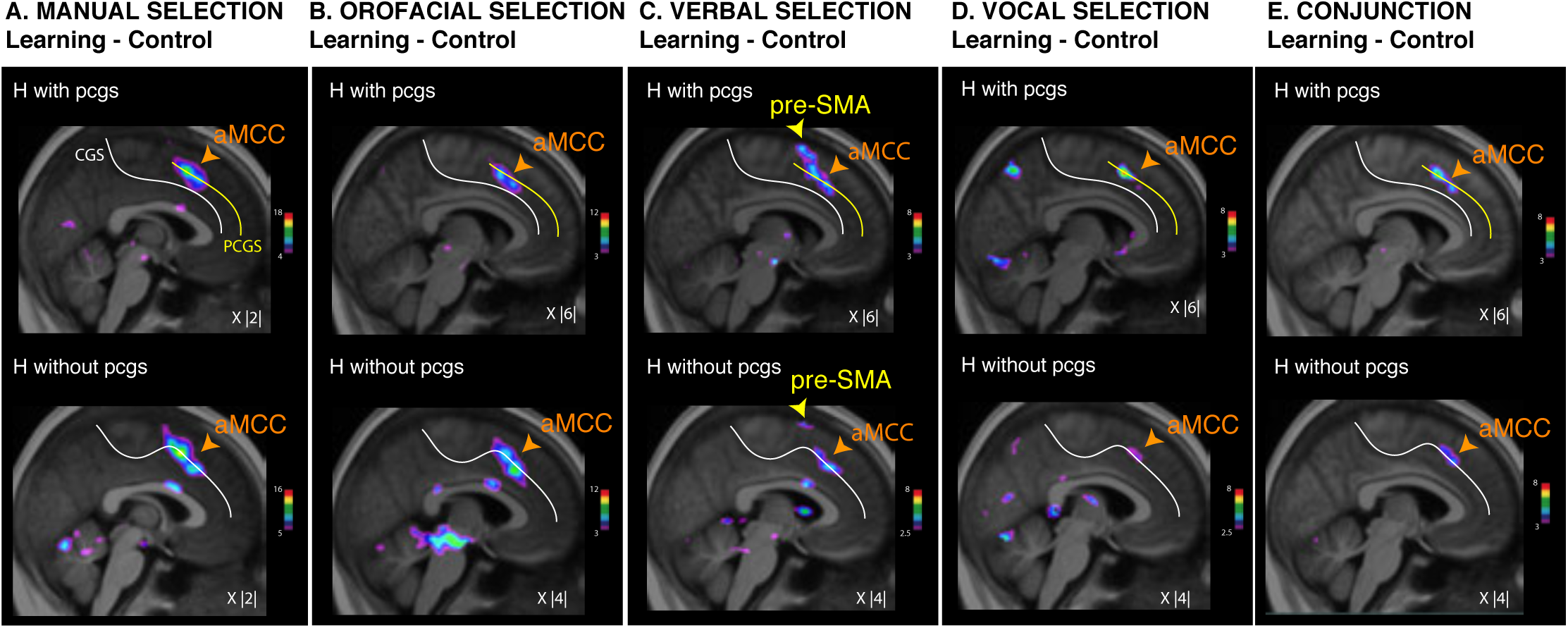
Conditional learning during response selection in the medial frontal cortex. Top panel: group analysis results of increased activity during response selection in the learning period in comparison with control trials in hemispheres displaying a cingulate (cgs) and a paracingulate sulcus (pcgs). Bottom panel: group analysis results of increased activity during response selection in the learning period in comparison with the control trials in hemispheres displaying no paracingulate sulcus. A. during manual response selection, B. during orofacial response selection, C. during verbal response selection, and D. during vocal/verbal response selection. E. Top panel: conjunction analysis between the 3 contrasts presented in the group analysis in A, B, C, and D in hemispheres displaying both a cgs and a pcgs in A, B, and C. Bottom panel: conjunction analysis between the 3 contrasts presented in hemispheres displaying no pcgs in A, B, C, and D in hemispheres displaying no pcgs. The color scales represent the range of the t-statistic values. The X values correspond to the medio-lateral level of the section in the MNI space.

In view of the significant inter-subject and inter-hemispheric sulcal variability observed in the medial frontal cortex ^30,31^, we performed sub-group analyses of the learning minus control contrast (during response selection) separately for hemispheres with a paracingulate sulcus (pcgs), and hemispheres without pcgs (Fig 4, Table S2). In both pcgs and no-pcgs groups, we observed two foci of increased activity in the anterior mid-cingulate cortex (aMCC) across all four response conditions (Fig 4A, 4B, 4C, 4D, see Table S2 for peak locations and t-values). As demonstrated by a conjunction analysis (Fig 4E), the two aMCC peaks occupy the same locations across response modalities. Importantly, our results also revealed that increased aMCC activity were consistently observed in the pcgs when present, and in the cgs when the pcgs was absent (see Fig 4). These results suggest that the aMCC is involved in the conditional selection of all effector types during learning when the learning is based on auditory vocal/verbal feedback.

By contrast, the pre-SMA showed increased response selection activity only during the learning of visuo-verbal associations (Fig 4C, Table S2), and not for learning associations between visual stimuli and the other response effectors (manual, orofacial, and nonspeech vocal). This point is confirmed in Fig 4E which shows that the pre-SMA does not display increased activity in the conjunction analysis across the various response conditions.

### The VLF-DMF network is involved in the analysis of auditory vocal/verbal feedback (FB) during conditional associative learning

To identify the brain regions associated with the analysis of auditory nonspeech vocal and verbal feedback during the learning of conditional relations, we contrasted, respectively, 1) the BOLD signal at the vocal feedback epochs in learning versus control trials with the same motor effector, and 2) the BOLD signal at the verbal feedback epochs in learning versus control trials with the same motor effector.

During the learning of visuo-manual associations, the analysis of nonspeech vocal and verbal feedback showed increased activity in ventral area 44 and the MCC (Fig 5A, Table S3). Additionally, we also observed increased activity in the PMd during the processing of both vocal and verbal feedback (Fig 5A, Table S3). This finding was congruent with previous work^25^ that revealed increased PMd activity as subjects processed visual behavioral feedback during visuo-manual associative learning.

**Figure 5.**
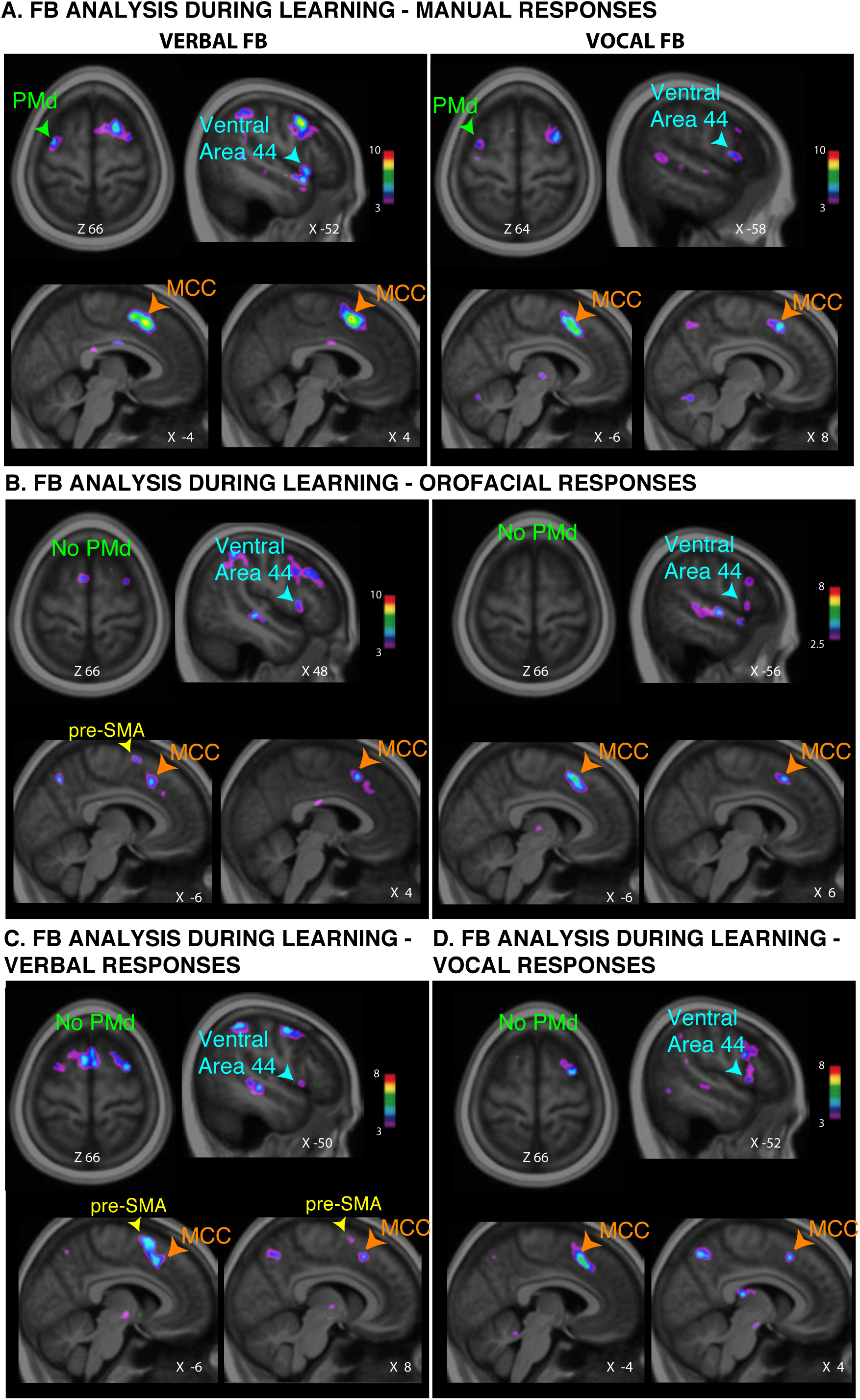
Verbal versus vocal feedback analysis during the visuo-motor conditional learning period in the posterior lateral prefrontal and medial frontal cortex. A. Group analysis results displaying increased activity during the analysis of verbal (left panel) and vocal (right panel) feedback during learning visuo-manual conditional associations compared to that during the analysis of verbal (left panel) and vocal (right panel) feedback during visuo-manual control trials. B. Group analysis results displaying Increased activity during the analysis of verbal (left panel) and vocal (right panel) feedback during learning visuo-orofacial conditional associations compared to that during the analysis of verbal (left panel) and vocal (right panel) feedback during visuo-orofacial control trials. C. Group analysis results displaying Increased activity during the analysis of verbal feedback during learning visuo-verbal conditional associations compared to that during the analysis of verbal feedback during visuo-verbal control trials. D. Group analysis results displaying Increased activity during the analysis of vocal feedback during learning visuo-vocal conditional associations compared to that during the analysis of vocal feedback during visuo-vocal control trials. The color scales represent the ranges of the t-statistic values. The X and Z values correspond to the medio-lateral and dorso-ventral levels of the section in the MNI space, respectively.

During the learning of visuo-orofacial associations, the analysis of nonspeech vocal and verbal feedback also showed increased activity in ventral area 44 and the MCC (Fig 5B, Table S3). In contrast to the visuo-manual condition, there was no increased activity in the PMd. Finally, there was increased activity in the pre-SMA only during the processing of verbal feedback, and not nonspeech vocal feedback (Fig 5B, Table S3).

During the learning of visuo-vocal/verbal associations, the analysis of nonspeech vocal and verbal feedback also showed increased activity in ventral area 44 and the MCC (Fig 5C, 5D; Table S3). In contrast to the visuo-manual condition, there was no increased activity in the PMd. Finally, there was increased activity in the pre-SMA only during the processing of verbal feedback and not nonspeech feedback (Fig 5C, 5D; Table S3).

In summary, during conditional associative learning, a common set of regions including ventral area 44 and the MCC is involved in the analysis of both vocal and verbal feedback to drive the learning of manual, orofacial, nonspeech vocal, and speech vocal conditional relations. The PMd region appears to be specifically recruited in feedback analysis for learning associations involving manual, but not orofacial and nonspeech and speech vocal responses. Interestingly, the pre-SMA appeared to be specifically recruited for the processing of verbal feedback for the learning of orofacial and nonspeech and speech vocal conditional associations. This suggests that the pre-SMA could have a particular role in exerting cognitive control on verbal and orofacial responses and to monitor performance based on verbal feedback specifically.

To identify the precise location of the MCC region involved in auditory verbal/vocal processing, we compared the BOLD signal during the analysis of vocal and verbal feedback in learning versus control trials, in hemispheres with and without a pcgs (see Methods). The results revealed that vocal and verbal FB processing are performed in the pcgs when present and in the cgs when the pcgs is absent (Fig 6). The same region is involved across the 3 effectors of responses (manual-Fig 6A-B, orofacial-Fig 6C-D, verbal-Fig 6E, vocal-Fig 6F) as shown by the conjunction analysis (Fig 6G).

**Figure 6.**
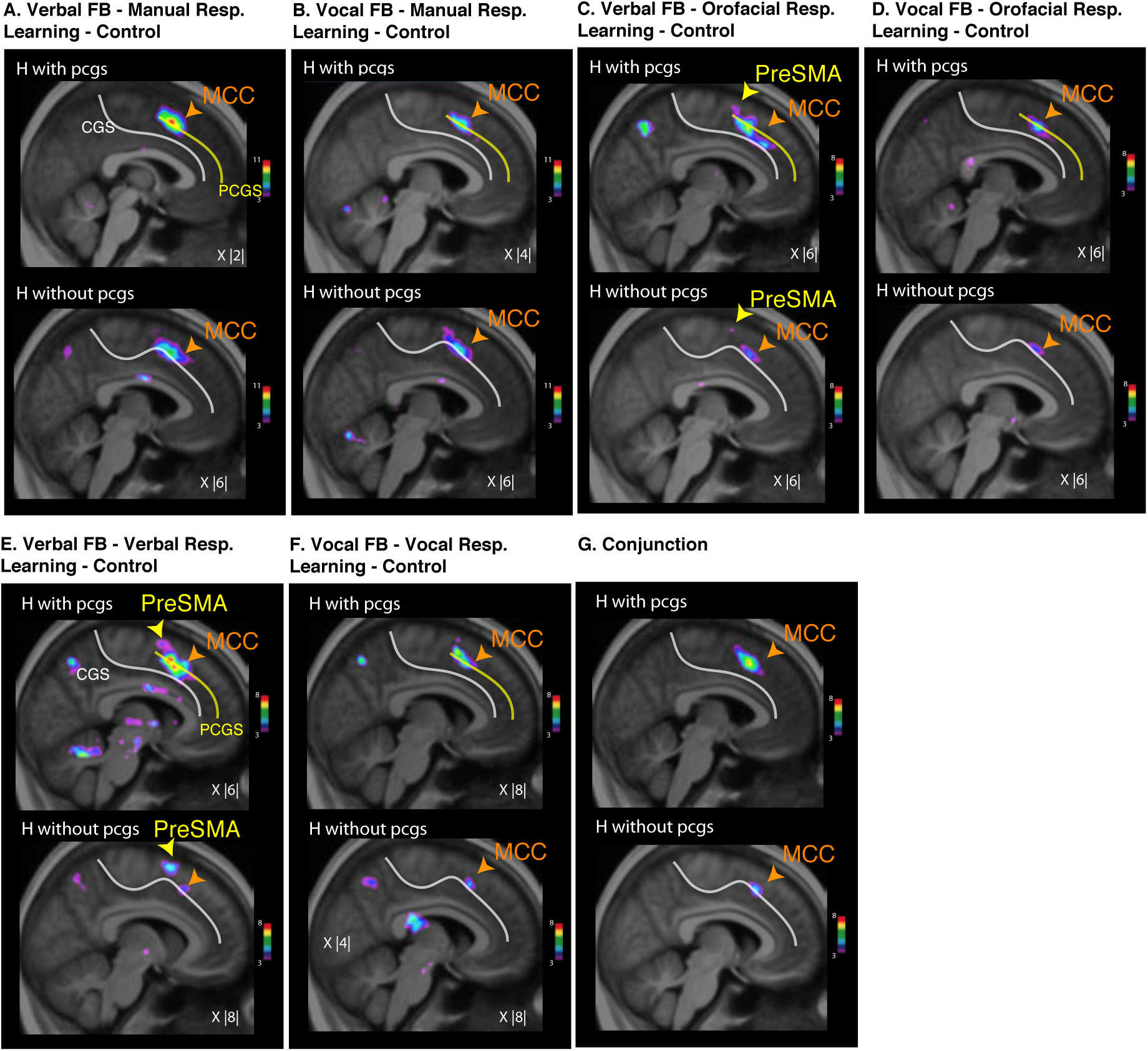
Group analysis of vocal/verbal feedback during visuo-motor conditional learning in the MCC in the group analysis (top panels), in hemispheres (H) exhibiting both a cingulate sulcus (cgs) and a paracingulate sulcus (pcgs) (middle panels) and in hemispheres without a paracingulate sulcus (bottom panels). A. Comparison between the BOLD signal at the occurrence of feedback during learning visuo-manual conditional associations versus during visuo-manual control trials. B. Comparison between the BOLD signal at the occurrence of feedback during learning visuo-orofacial conditional associations versus during visuo-orofacial control trials. C. Comparison between the BOLD signal at the occurrence of feedback during learning visuo-vocal/verbal conditional associations versus during visuo-vocal/verbal control trials. D. Conjunction between the contrasts presented in A, B, and C in hemispheres displaying or not a pcgs. The color scales represent the ranges of the t-statistic values. The X values correspond to the medio-lateral levels of the section in the MNI space, respectively.

### The MCC region that is involved in the analysis of auditory vocal/verbal feedback is the face motor representation of the cingulate motor area

To identify more precisely the brain regions associated with the analysis of vocal/verbal feedback during learning, we compared the BOLD activity at the occurrence of feedback during learning versus during post-learning trials. This specific contrast was used to relate findings to previous studies assessing the neural basis of feedback analysis in tasks including a learning and a post-learning period ^28^. We assessed the six response-feedback combinations: Manual-Vocal, Manual-Verbal, Vocal-Vocal, Verbal-Verbal, Orofacial-Vocal and Orofacial-Verbal (Fig 7). This analysis showed that the increased activities were consistently found in the same MCC region across the three response modalities (manual – red squares/circles, vocal-verbal – dark blue squares/circles, orofacial – green squares/circles) and two feedback types (vocal – circles, verbal squares). Specifically, individual increased activities were found close to the intersection between the posterior vertical paracingulate sulcus (p-vpcgs) with the cgs when there is no pcgs and with the pcgs when present. This position likely corresponded to the location of the face motor representation of the anterior rostral cingulate zone (RCZa) ^21^ (Fig 7). To confirm if these vocal feedback processing-related activation peaks corresponded to the RCZa ‘face’ motor representation, we compared their locations with the average activation peaks corresponding to the tongue and eye RCZa motor representations obtained in a previous study from the same set of subjects ^33^. As displayed in Fig 7, the feedback-related peaks obtained in the current study were located near the average RCZa face (eye and tongue) representation.

**Figure 7.**
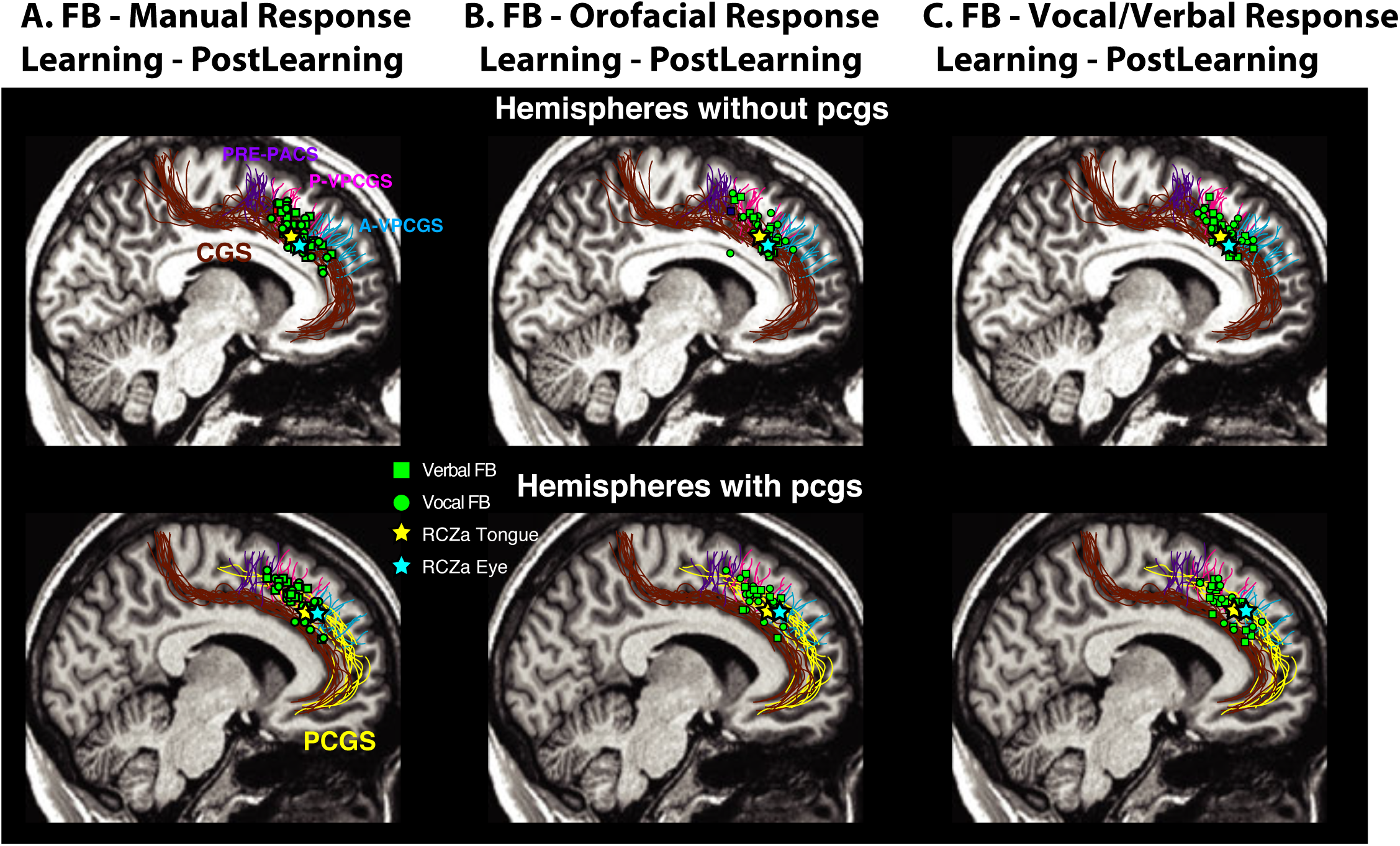
Auditory vocal/verbal feedback processing during learning visuo-motor conditional associations in relation to the anterior cingulate motor area. A. Comparison between the BOLD signal at the occurrence of verbal (squares) and vocal (circles) feedback during learning visuo-manual conditional associations versus during visuo-manual control trials in individual subjects (each square/circle represent the location of increased activity of one subject) and in hemispheres displaying both a cingulate sulcus (cgs) and a paracingulate sulcus (pcgs, top diagrams) and in hemispheres without a pcgs (bottom diagrams). B. Identical as A but increased activities are based on the comparison between the BOLD signal at the occurrence of feedback during learning visuo-orofacial conditional associations versus during visuo-orofacial control trials. C. Identical as A and B but increased activities are based on the comparison between the BOLD signal at the occurrence of feedback during learning visuo-vocal/verbal conditional associations versus during visuo-verbal/vocal control trials. Yellow and blue stars represent average locations of the tongue and eye motor representations in the RCZa that were derived from a motor mapping task in Loh et al. 2018. Abbreviations: cs, cingulate sulcus; pcgs, paracingulate sulcus; pre-pacs, pre-paracentral sulcus; a/p-vpcgs, anterior/posterior vertical paracingulate sulcus.

## Discussion

The present study is designed to reveal the roles of the various ventrolateral-dorsomedial frontal (VLF-DMF) cortical regions in human cognitive vocal and orofacial control that might be generic across primates, and the potential changes that could have occurred in this network across primate evolution to enable speech production. Providing critical insights into the above question, our key results have demonstrated important functional dissociations between the various components of the human VLF and DMF regions in the cognitive control of orofacial and vocal actions.

Firstly, within the VLF, the ventral area 44 was specifically involved in the cognitive conditional selection between competing orofacial, nonverbal vocal and verbal vocal acts during both the acquisition and execution of such responses. Importantly, it was not involved during the learning and execution of conditional <u>manual</u> response selection, which, instead, recruited the dorsal premotor cortex (PMd). Single-subject analyses confirmed that ventral area 44 peaks were consistently situated in the pars opercularis, i.e. the region bordered by the inferior frontal sulcus, the Sylvian fissure, the ascending ramus of the Sylvian fissure and the inferior precentral sulcus. Thus, the results support the hypothesis that primate area 44 plays a critical role in the cognitive selection of orofacial and nonverbal and verbal vocal responses ^2,34^. This hypothesis was based on the anatomical connectivity profile of area 44, namely the fact that it is located immediately in front of the precentral gyrus orofacial region and is directly connected to it ^35^. Frey and colleagues ^36^ have shown that area 44 has rich links with the prefrontal cortex (area 45 and mid-dorsolateral area 9/46V), receives somatosensory inputs from the parietal operculum, insula and the inferior parietal cortex, and has strong connections with both lateral and medial premotor regions (ventral premotor area, cingulate areas 23 and 24, supplementary motor areas). This pattern of connectivity indicates that area 44 is well-placed to mediate top-down cognitive (prefrontal cortex) influences on vocal and orofacial motor acts (premotor areas). Moreover, several functional neuroimaging studies have also implicated area 44 in speech control and production ^37,38^. Indicative that the role of ventral area 44 in cognitive orofacial and vocal selection could be conserved across primates, the same region in monkeys is also associated with vocal and orofacial control: electrical stimulation in the ventral extent of the fundus of the inferior arcuate sulcus, where the macaque area 44 is found, has been shown to evoke orofacial movements ^15^. Neural activity in this area has been associated with the conditioned production of calls ^3,39^, suggesting its role in volitional vocal control.

The ventral area 44 is also involved in the analysis of vocal nonverbal and verbal feedback during conditional associative learning across all response modalities. This observation is congruent with both human and monkey studies that found that the ventrolateral prefrontal cortex (vlPFC) is not only involved in the production of vocal responses but also in the processing of auditory information during vocal adjustments. In monkeys, Hage and Nieder ^40^ found that the same vlPFC neurons that are involved in conditioned vocal productions also responded to auditory information. They suggested that this mechanism could underlie the ability of monkeys to adjust vocalizations in response to environmental noise or calls by conspecifics. In human subjects, Chang et al. ^41^ have shown, via intracortical recordings, that as subjects adjusted their vocal productions in response to acoustic perturbations, the ventral prefrontal cortex reflected compensatory activity changes that were correlated with both the activity associated with auditory processing and the magnitude of the vocal pitch adjustment. Functional neuroimaging investigations have also shown that human area 44 shows increased activity during both the processing of articulatory/phonological information and the production of verbal responses ^32^. As such, the observed role of the human ventral area 44 in the cognitive processing of auditory vocal/verbal information could also be preserved across primates.

Importantly, the present study demonstrated a major difference between activations of area 44 and area 45, although both these regions are usually considered to be part of Broca’s region in the language dominant hemisphere ^42^. Unlike area 44, which was involved in orofacial, vocal nonverbal and verbal response selections both during the learning and the post-learning period, activation in area 45 was related only to the conditional selection of nonverbal vocal and verbal responses, and only during the learning period. Thus, the dysgranular transitional area 44 that lies between the ventral premotor cortex that controls the orofacial musculature, and the cognitive prefrontal area 45, appears to be the fundamental area regulating orofacial/vocal output selections regardless of whether these selections are just orofacial movements, nonspeech vocal or speech vocal responses and whether these selections occur during the learning or post-learning periods. By contrast, area 45 contributes only to nonverbal vocal and speech selections and only during the learning period, namely the period when the conditional relations are not well learned and the subject must therefore engage in active mnemonic retrieval. This finding is consistent with the hypothesis that area 45 is critical for controlled memory retrieval ^8,34,35^. There is functional neuroimaging evidence of the involvement of area 45 in the controlled effortful mnemonic retrieval of verbal information, such as the free recall of words that appear within particular contexts ^9^ and a more recent study has shown that patients with lesions to the ventrolateral prefrontal region, but not those with lesions involving the dorsolateral prefrontal region, show impairments in the active controlled retrieval of the contexts within which words were presented ^43^. The above findings regarding the differential involvement of the two cytoarchitectonic areas that comprise Broca’s region are consistent with the hypothesis that the prefrontal granular component, i.e. area 45, is the critical element for the cognitive controlled retrieval of verbal information, which is then turned into speech utterances by the adjacent dysgranular area 44, leading to the final motor output via the precentral orofacial region ^34^. With its specific role in verbal retrieval, area 45 could be considered the part of the VLF that has changed in humans to support speech production. In monkeys, area 45 has been found to contain neurons that respond to both species-specific vocalizations and faces, and receives strong audio-visual inputs from the temporal cortices, indicating a role in the active retrieval and integration of audio-visual information ^44^. As such, from monkeys to humans, area 45 might have evolved from the retrieval and integration of basic audio-visual communicative information (faces and vocalisations) to the more complex multimodal inputs that are inherent in speech ^40^.

Lastly, our results also revealed a novel dorso-ventral functional dissociation within the pars opercularis (area 44) itself. Distinct from the ventral area 44 discussed above, dorsal area 44 was involved specifically during the learning period of visuo-motor conditional associations across all response modalities, and not during the execution of learnt associations. In support of this dissociation, recent parcellations of the pars opercularis on the basis of cytoarchitecture ^45^, receptor architecture ^46^, and connectivity ^47^ have demonstrated that area 44 can be further sub-divided into dorsal and ventral parts. Recent neuroimaging studies have also shown functional dissociations between the dorsal and ventral area 44. However, the precise roles attributed to the two sub-regions are currently still debated. For instance, Molnar-Szakacs and colleagues ^48^ found that the dorsal area 44 was activated during both action observation and imitation, while the ventral area 44 was activated only during action imitation. In agreement, Binkofski et al. ^36^ also showed that the ventral, but not dorsal area 44, was implicated during movement imagery. Lastly, during language production, the ventral area 44 was found to be involved in syntactic processing ^50^ and comprehension ^51^, while the dorsal area 44 was involved in phonological processing ^52^. As such, our findings are clearly in agreement with the emerging view that the pars opercularis can be sub-divided into dorsal and ventral parts.

With regards to the DMF, our findings indicated that the MCC was involved during the learning of conditional responses based on auditory vocal and verbal feedback. Note that the MCC was <u>not</u> involved in response selection during the post-learning period, indicating its specific role in adaptive learning, in agreement with previous studies ^25,26,28^. Furthermore, the role of the MCC during learning visuo-motor conditional associations is not effector-specific as the same MCC region is activated during visuo-manual, orofacial and nonverbal vocal and verbal conditional associative learning. Importantly, subject-by-subject analyses further indicated that the activation focus in the MCC corresponds to the ‘face’ motor representation within the anterior mid-cingulate motor region (RCZa). In other words, nonverbal vocal and verbal feedback processing was carried out in the ‘face’ motor region in the RCZa. Our findings indicate that the ‘face’ motor representation of RCZa, within the MCC, contributes to the processing of auditory vocal and verbal feedback for behavioral action adaptation. Consistent with our results, accumulating evidence from both monkey and human functional investigations converges on the role of the primate MCC in driving behavioral adaptations via the evaluation of action outcomes ^25,29,53–56^. Importantly, based on a review of the locations of outcome-related and motor-related activity in the monkey and human MCC, Procyk and colleagues ^53^ reported an overlap between the locations for the evaluation of juice-rewarded behavioral outcomes and the face motor representation in the monkey rostralmost cingulate motor area (CMAr), strongly suggesting that behavioral feedback evaluation in the MCC is embodied in the CMAr motor representation corresponding to the modality of the feedback. The present study supports this hypothesis showing that adaptive auditory feedback is being processed by the face motor representation in the human homologue of the monkey CMAr, i.e. RCZa.

The preSMA is selectively recruited during the learning of conditional verbal (but not nonverbal vocal, orofacial, or manual) response selections based on verbal (but not nonverbal vocal) feedback. These findings highlight the special role of preSMA in the learning of verbal responses and the processing of verbal feedback for such learning. The current literature suggests that the preSMA is involved in the learning of associations between visual stimuli and sequences of motor actions ^57,58^. A possible explanation of the preSMA’s unique involvement in the learning of visuo-verbal associations in the current study might be that only verbal responses involve a sequence of motor acts (i.e. sounds), whereas manual (single button presses), orofacial (single mouth movements) and vocal responses (single sounds) involve individual motor actions. In support of the preSMA’s role in verbal processing, Lima and colleagues ^59^ have shown that the preSMA is often engaged in the auditory processing of speech. Importantly, these investigators also suggested that the preSMA is involved in the volitional activation/retrieval of the specific speech motor representations associated with the perceived speech sounds. This could explain our observation that the preSMA was active during both the processing and selection of verbal responses during learning. The role of the preSMA in the learning of context-motor sequence associations that is observed in the macaque ^60^ appears to be conserved in the human brain. Although nonhuman primates do not produce speech, it has been shown that the preSMA in monkeys is associated with volitional vocal production ^2^: Stimulation in the pre-SMA produces orofacial movements ^17^. Lesions of the preSMA region lead to increased latencies of spontaneous and conditioned call productions ^61^. Based on these findings, it appears that the role of preSMA in the volitional control of orofacial/vocal patterns may have been adapted in the human brain for the control of speech patterns via context-motor sequence associations.

Together, our results demonstrate that within the human VLF-DMF network, ventral area 44 and MCC could have conserved functions in primate cognitive vocal control: while ventral area 44 is involved in the cognitive selection of vocal and orofacial actions, as well as the active processing of auditory-vocal information, the MCC is involved in the evaluation of vocal/verbal feedback that leads to behavioural adaptation in learning conditional associations between vocal/orofacial actions and arbitrary external visual stimuli. Indeed in a previous review ^2^, we have argued that the above functional contributions of area 44 and MCC are generic across primates based on anatomical and functional homologies of these regions in cognitive vocal control. Within the human VLF- DMF, area 45 and the preSMA could be regions that are specialized for speech production: Area 45 is recruited for the selective retrieval of verbal/semantic information that will be turned into orofacial action by area 44, while the preSMA is specifically involved in driving verbal action selections based on auditory verbal feedback processing.

Apart from these specializations within the VLF-DMF network, another important change that could have contributed to the emergence of human speech capacities is the emergence of a cortical laryngeal representation in the human primary motor orofacial region that afforded increased access to fine-motor control over oro-laryngeal movements^62^. As such, ventral area 44, with strong connections to the primary motor face area via the ventral premotor area, would be able to exercise control via conditional sensory-vocal associations over a wider range of oro-laryngeal actions. The preSMA, which is also strongly linked to the primary motor face representation, via the SMA, would also be able to build context-motor sequence associations with complex speech motor patterns and activate them based on their auditory representations. The MCC, which is directly connected to the ventral premotor area, would also be able to drive oro-laryngeal adaptations, based on feedback evaluation, at the fine motor level. Finally, area 45, would provide semantic and other information retrieved from lateral temporal cortex and posterior parietal cortex that would bring the VLF-DMF network in the service of higher cognition in the language dominant hemisphere of the human brain ^34,42^. This could explain the expanded capacity of the human brain to generate flexibly and modify vocal patterns.

## Methods

### Subjects

22 healthy, right-handed native French speakers were recruited to participate in a training session and 3 fMRI sessions. Data from two subjects (S2, S13) were omitted from the analyses because they had shown poor performance across the three experimental sessions. Two other subjects (S6, S9) did not participate in any of the sessions because of claustrophobia. Consequently, the final dataset consisted of 18 subjects (10 males, mean age=26.22 (SD=3.12)). The study was carried out in accordance with the recommendations of the Code de la Santé Publique and was approved by the “Agence Nationale de Sécurité des médicaments et des produits de santé (ansm)” and the “Comité de Protection des Personnes (CPP) Sud-Est III” (N° EudraCT: 2014-A01125-42). It also received a Clinical Trial Number (NCT03124173, see https://clinicaltrials.gov). All subjects provided written informed consent in accordance with the Declaration of Helsinki.

### Behavioral tasks

Subjects performed two tasks: 1) A visuo-motor conditional learning task (Fig. 1A), and 2) a visuo-motor control task (Fig. 1B). The sequence of events was comparable for the two tasks: an instruction screen lasting for 2s informed the participant of the type of task to be performed.: “Find the correct associations” (which indicated the selection of the response linked to the particular cue presented on each trial) for the conditional learning task or “Select the X response” (which indicated the particular response to be performed on each trial) for the control task block, respectively. Following the instructions, a black screen with a central fixation cross was presented during a jittered inter-trial interval of 0.5-8s (mean=3.5s). One of three possible visual stimuli (abstract grayscale images) was then presented on the center of the screen for 2 seconds during which the subject had to select and perform one of three possible motor responses (learning task) or perform an instructed motor response (control task). Stimulus presentation was ordered as randomly permuted blocks of the three possible images (i.e. abc, bca, acb, etc.). After a jittered delay (0.5-6s, mean=2s), a vocal or verbal feedback (1s) was provided to inform the subject whether the response selected and performed was the correct one for the presented image (learning task) or the correct instructed response (control task). After a jittered inter-trial interval (0.5-8s, mean=3.5s), another trial started with the presentation of one of the 3 possible stimuli. In half of the task blocks, verbal feedback was provided (positive: “Correct”; negative: “Error”). For the remaining blocks, vocal feedback was provided (positive: “Aha”; negative: “Boo”). The subject was informed about the type of feedback that would be provided after the response performance by the text color on the instruction screen at the start of each task (Verbal: Yellow, Vocal: Red). The four different types of vocal/verbal feedback were recorded from the same male voice and processed using Audacity software.

In each visuo-motor conditional learning task block (Fig. 1A), subjects had to acquire three visuo-motor conditional associations (if visual stimulus A, then motor response X, if visual stimulus B, then motor response Y, and if visual stimulus C, then motor response Z) via trial-and-error. On every trial, one of the three possible stimuli were presented and the subject had to select one of the three possible responses to perform (Response Selection). Auditory vocal/verbal feedback was then provided to inform the subject whether the selected response was the correct one for the presented stimulus. This trial-and-error learning period continued until the subject identified the correct response for each stimulus. When a correct response had been performed once for each of the three visual stimuli, the task proceeded to the “post-learning” period during which the subject had to repeat, in response to the appropriate cues, each of the learnt conditional associations twice (6 trials). Note that in each visuo-motor conditional learning task block, a novel set of 3 stimuli was presented and, therefore, the subject had to learn the new stimulus-response relations.

In each visuo-motor control task block (Fig. 1C), subjects were informed of the specific response to perform on the instruction screen. Subsequently, they had to perform that particular instructed response to all of the three different visual stimuli over five trials. Thus, by contrast to the learning task, during the control task the subjects did not have to select the appropriate motor response to perform based on learning of the correct stimulus to response arbitrary relations nor to adjust their selections based on the provided feedback. Different sets of abstract visual images were used in all learning and control task blocks. Note that in each control task block, a novel set of 3 stimuli was presented.

In the present study, there were three versions of the visuo-motor conditional learning and control tasks that corresponded to three different response effectors: manual, orofacial, vocal/verbal (Fig. 1B). In the visuo-manual condition, the subjects acquired associations between three different hand presses on an MRI compatible button box (Current Designs, Philadelphia, USA) and visual stimuli in the conditional learning task and performed instructed button hand presses in the control task.

In the visuo-orofacial condition, subjects performed the conditional learning task and control tasks using three different orofacial movements (Fig 1B, middle panel). In the visuo-vocal/verbal condition (Fig 1B, right-most panel), the responses were either three different meaningless vocal responses (“Ahh”, “Ihh” and “Ohh”) or verbal responses (the French words “Bac”, “Vis” and “Col” during the learning and control tasks and the feedback provided was either nonverbal vocal or verbal. These nonspeech vocal and verbal responses were selected to match, as closely as possible, the orofacial movements performed in the visuo-orofacial condition: the first orofacial action (top image in the orofacial panel; Fig 1B) is almost identical to the mouth movements engaged in producing the vocal action ‘AAH’ and similar to the verbal action “BAC”. In the same manner, the second and third orofacial actions corresponded to vocal actions “IIH” and “OHH”, and verbal actions “VIS” and “COL”, respectively. Subjects were informed of which set of responses to use via the text color of the instructions (red – “vocal”, yellow - “verbal”). For the manual and orofacial conditions, subjects used the same set of responses for verbal and vocal feedback.

Each subject participated in three fMRI sessions. In the first session (visuo-manual fMRI session), they performed the visuo-manual conditional learning task and the visuo-manual control task. In the second session (visuo-orofacial fMRI session), they performed the visuo-orofacial conditional learning task and the visuo-orofacial control task. In the third session (visuo-vocal/verbal fMRI session), they performed the visuo-vocal/verbal conditional learning task and the visuo-vocal/verbal control task. To ensure optimal performance during the actual fMRI sessions, all subjects were familiarized with all three versions of the learning and control tasks in a separate training session held outside the scanner. During the training session, subjects practiced the visuo-manual, visuo-orofacial, and visuo-vocal/verbal conditional learning tasks until they consistently met the following criteria in each version: 1) not more than one sub-optimal search (i.e. trying the same incorrect response to a particular stimulus or trying a response that had already been correctly associated to another stimulus) during the learning phase, and 2) not more than one error in the post-learning phase.

### Behavioral data acquisition

In the visuo-manual fMRI session, subjects responded directly to the task within the scanner via an MRI-compatible button box (Current Designs, Philadelphia, USA) connected to the Presentation computer. Raw trial data, including trial events, reaction times and percent correct responses were directly extracted from the Presentation logfiles.

In the visuo-vocal/verbal fMRI session, an MRI-compatible microphone (MO 2000 model, Seinnheiser Electronic, Germany) was installed on the head-coil, near the subjects’ mouths, to record their vocal/verbal responses during the task. Outside the scanner, an experimenter monitored the subjects’ vocal/verbal responses in real-time and responded to the task via keyboard presses on the Presentation computer. A Biopac MP150 system (Biopac Systems Inc, Goleta, CA) was used to acquire simultaneously, and synchronize: 1) the analog audio signals from the microphone, 2) task event signals from Presentation, and 3) TTL signals from the MRI scanner. After the experiment, the subjects’ actual vocal/verbal response onsets were computed from the analog audio signal (time-synchronized to task events and MRI pulses) using a customized onset detection algorithm (written by R. Neveu and available upon request) implemented on Matlab (www.mathworks.com). The detected vocal/verbal responses in the audio recordings were subsequently used to verify the responses made by the experimenter during the task (logged by the Presentation software). The trial event timings were acquired directly from the Presentation logfiles.

In the visuo-orofacial fMRI session, a home-made video-camera was used to record the subjects’ mouth responses as they performed the task in the scanner. The camera was installed outside the scanner tunnel and positioned to provide a view of the subjects’ mouth from a mirror fixed on the head coil. As in the visuo-vocal/verbal experiment, an experimenter responded to the task outside the scanner according to the subjects’ mouth responses. The Biopac system was also used to acquire the video data, in parallel with task event and MRI TTL signals. A customised Matlab program (written by K. Loh and available upon request) was used to synchronize the video recordings with the timings of the task events, and subsequently, allow the experimenter to manually playback each video segment frame-by-frame to determine the onset of each orofacial response. The orofacial responses detected from the video were cross-checked with the responses made by the experimenter during the task. The trial event onsets were acquired directly from the Presentation logfiles.

### MRI acquisition

Scanning was performed on a 3T Siemens Magnetom Prisma MRI Scanner (Siemens Healthcare, Erlangen, Germany). To minimize movements during the motor tasks, the subjects’ heads were tightly cushioned throughout the acquisition. The main experimental protocol consisted of three MRI sessions (∼2hrs), each involving a different version of the visuo-motor conditional learning and control tasks (visuo-manual, visuo-orofacial, and visuo-vocal/verbal). For each fMRI session, subjects performed four to six experimental runs in the scanner. Experimental runs were programmed and presented using Presentation (Neurobehavioral systems). Visual stimuli were presented via an LCD projector with a mirror system. Auditory feedback was delivered via MRI-compatible earphones (Siemens Healthcare, Erlangen, Germany).

The start of each run was synchronized to the 5th TTL pulse from the MRI scanner following scan initiation. Each run began with a fixation task followed by two motor mapping task blocks involving either: hand and eye movements (visuo-manual session), verbal and nonspeech vocal responses (visuo-vocal/verbal session) or mouth and tongue movements (visuo-orofacial session). The details of the fixation and motor mapping tasks are described in Loh et al. ^33^. After the three fixation/motor mapping task blocks (total duration = 28.5s), subjects were presented with the six learning task blocks (3 x 2 feedback types) and six control task blocks (3 motor responses x 2 feedback types) that were randomly interleaved in each run. The total length of each run was limited to a maximum of 14.6min, yielding 400 T2*-weighted gradient echo planar EPI volumes (40 descending oblique slices, voxel resolution = 2.7mm x 2.7mm x 2.7mm, TR = 2.2s, TE = 30.0s, flip angle = 90°). The initial five volumes of each acquisition were discarded to avoid confounds of unsteady magnetization. High-resolution T1 structural images (MPRAGE, 0.9mm^3^ isotropic voxels, 192 slices, TR=3.5s, TE=2.67s), together with Diffusion Tensor Imaging (DTI) and resting state functional scans (not analyzed in the current study) were each acquired in one of the three MRI sessions at the end of all experiment task runs.

### MRI analyses

The preprocessing and analyses of MRI data were performed with Statistical Parametric Mapping software (SPM12; Wellcome Department of Cognitive Neurology, University of College London, London, UK; http://www.fil.ion.ucl.ac.uk/spm) and Matlab 15b (www.mathworks.com).

First, structural and functional images were reoriented by setting their origins to the anterior commissure. The first 5 volumes of each run were excluded to remove T1 equilibrium effects. Next, within each session, we realigned all successive images to the first image of the session. Using the Artefact Detection Toolbox (ART; http://www.nitrc.org/projects/artifact_detect/), motion outliers were computed from the realigned images and realignment parameters. The realignment parameters and detected motion outliers were saved as covariates to model potential nonlinear head motion artefacts in subsequent statistical analyses. Slice-timing correction was applied with the time centre of the volume as reference. The subject-mean functional images were co-registered with the corresponding structural images using mutual information optimization. Functional and structural images were then spatially normalized into standard MNI space. Finally, functional images were smoothed using a 6-mm full-width half-maximum Gaussian kernel ^63–65^.

For each subject, fMRI data from the 3 fMRI sessions (manual, orofacial, and nonspeech vocal/verbal) were modeled separately. At the first level, each trial was modeled with impulse regressors at the two main events-of-interest: 1) Response Selection (RS) – the 2s epoch after the stimulus onset, during which the subject had to perform a response after stimulus presentation, and 2) Auditory Feedback (FB) – the 1s epoch after the onset of auditory feedback. RS and FB epochs were categorized into either learning (RS_L_, FB_L_), post-learning (RS_PL_, FB_PL_) or control (RS_C_, FB_C_) trial events. These regressors were then convolved with the canonical hemodynamic response function and entered into a general linear model of each subject’s fMRI data. The six scan-to-scan motion parameters produced during realignment and the ART-detected motion outliers were included as additional regressors in the general linear model to account for residual effects of subject movement.

To assess the brain regions involved in the visuo-motor conditional response selection, we contrasted the blood oxygenation-level dependent (BOLD) signal during RS_L_ and RS_PL_ events, where subjects actively selected their responses on the basis of the presented stimulus, with RS_C_ events, where subjects performed instructed responses. The two main contrasts (i.e. RS_L_ - RS_C_ and RS_PL_ - RS_C_) were examined for each response version at the group level and at the subject-by-subject level. At the group level, verbal and nonspeech vocal response selection trials are pooled in order to increase statistical power. To examine possible differences between nonspeech vocal and verbal responses, we distinguished between the two conditions in our subject-level analyses.

To determine the brain regions involved in the processing of auditory feedback during the learning of visuo-manual, visuo-orofacial, and visuo-vocal/visuo-verbal conditional associations, we examined the contrasts between FB_L_ and FB_PL_ events and between RS_L_ and RS_PL_ in each response version to determine if distinctive areas are involved in the processing of auditory feedback during the different response modalities. To determine whether verbal and nonspeech vocal feedback processing recruited different brain regions, we performed the above analyses separately for each response type (manual, nonverbal and verbal vocal, orofacial) with verbal or nonspeech vocal feedback.

Because of individual variations in cortical sulcal morphology in the dorsal premotor region (PMd), the ventrolateral Broca’s region and the medial frontal region, the above analyses were also assessed at the subject-by-subject level. In PMd, we identified activation peaks in relation to the dorsal branch of the superior precentral sulcus, the ventral branch of the superior precentral sulcus, and the superior frontal sulcus ^24,66^ (Fig 2). We identified activation peaks in relation to the limiting sulci of Broca’s region, i.e. the inferior precentral sulcus (iprs), the ascending ramus of the lateral fissure (aalf), the horizontal ramus of the lateral fissure (half), and the inferior frontal sulcus (ifs). In the medial frontal cortex, we identified activation peaks in relation to the cingulate sulcus (cgs), paracingulate sulcus (pcgs), and the vertical sulci joining the cgs and/or pcgs (i.e. the paracentral sulcus (pacs), the pre-paracentral sulcus (prpacs), and the posterior vertical paracingulate sulcus (p- vpcgs). It should be noted that the pcgs is present in 70% of subjects at least in one hemisphere and several studies have shown that the functional organization in the cingulate cortex depends on the sulcal pattern morphology. We, therefore, also performed sub-group analyses of fMRI data in which we separated hemispheres with a pcgs from hemispheres without a pcgs (see Amiez et al. ^28^ for the full description of the method).

**Figure 2.**
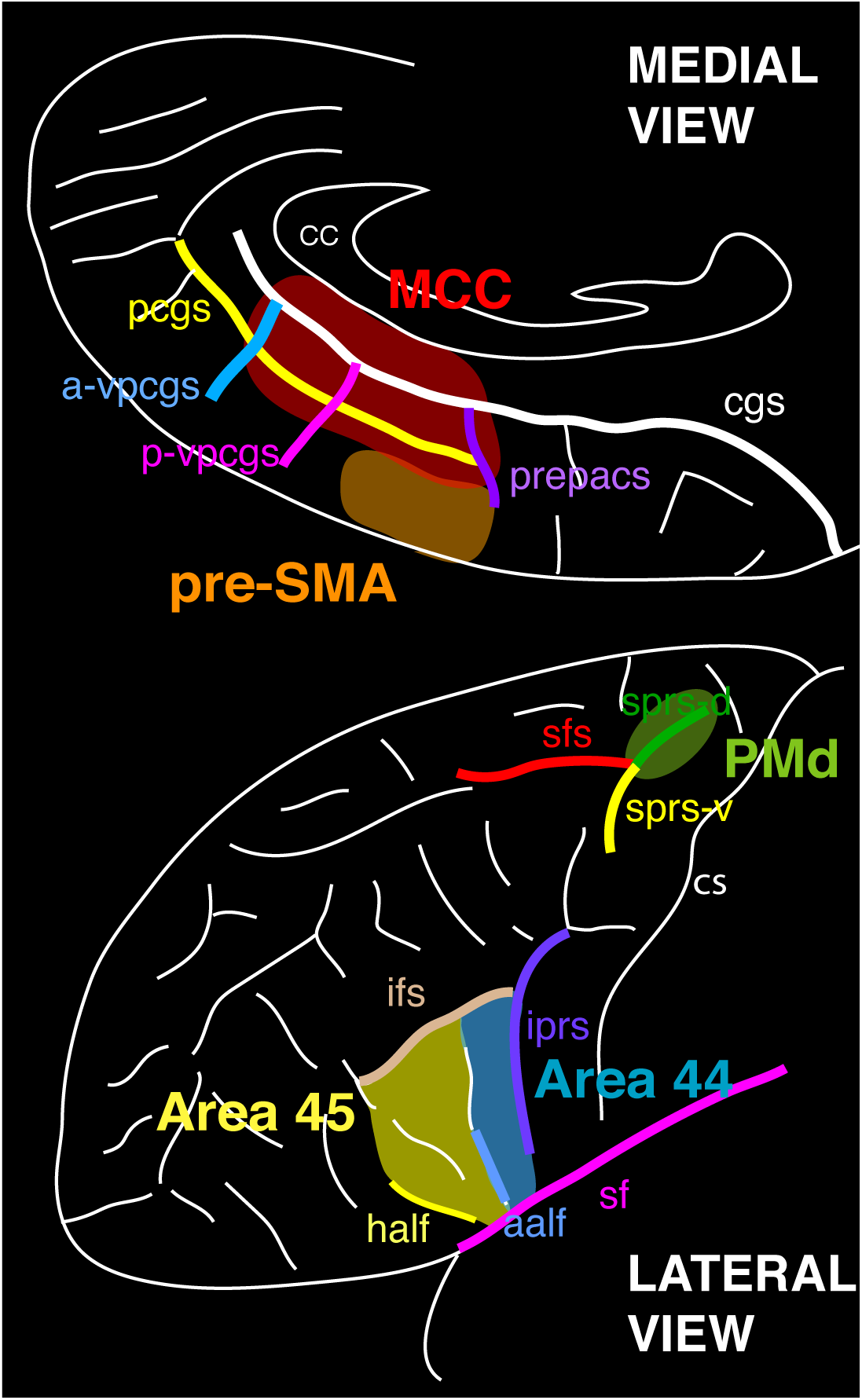
Sulci characteristic of the MCC and the preSMA (medial view), as well as areas 44 and 45 in Broca’s region and the dorsal premotor cortex (PMd) (lateral view). In the PMd the characteristic sulci are the dorsal branch of the superior precentral sulcus (sps-d), the ventral branch of the superior precentral sulcus (sps-v), and the superior frontal sulcus (sfs). Area 44 in Broca’s region is bounded by the inferior precentral sulcus (ips), the anterior ramus of the lateral fissure (aalf), and the inferior frontal sulcus (ifs). In the medial frontal cortex, the characteristic sulci are the cingulate sulcus (cgs), the paracingulate sulcus (pcgs), and the vertical sulci joining the cgs and/or pcgs, i.e. the paracentral sulcus (pacs), the pre-paracentral sulcus (prepacs), the posterior vertical paracingulate sulcus (p-vpcgs), and the anterior vertical paracingulate sulcus (a-vpcgs).

For the group, sub-group, and individual subject analyses, the resulting *t* statistic images were thresholded using the minimum given by a Bonferroni correction and random field theory to account for multiple comparisons. Statistical significance for the group analyses was assessed based on peak thresholds in exploratory and directed search, and the spatial extent of consecutive voxels. For a single voxel in a directed search, involving all peaks within an estimated grey matter of 600 cm^3^ covered by the slices, the threshold for significance (*p* < 0.05) was set at *t* = 5.18. For a single voxel in an exploratory search, involving all peaks within an estimated grey matter of 600 cm^3^ covered by the slices, the threshold for reporting a peak as significant (*p* < 0.05) was *t* = 6.77 ^67^. A predicted cluster of voxels with a volume extent >118.72 mm^3^, with a *t*-value > 3 was significant (*p* < 0.05) corrected for multiple comparisons ^67^. Statistical significance for individual subject analyses was assessed based on the spatial extent of consecutive voxels. A cluster volume extent > 444 mm^3^, associated with a *t* value > 2, was significant (*p* < 0.05), corrected for multiple comparisons ^67^.

## Data availability

The data and analysis scripts that support the findings of this study are available from the corresponding author upon request.

## Acknowledgments

This work was supported by the Human Frontier Science Program (RGP0044/2014), the Medical Research Foundation (FRM), the Neurodis Foundation, the French National Research Agency, CIHR Foundation FDN-143212, and by the labex CORTEX ANR-11-LABX- 0042 of Université de Lyon. K.L. was supported by the LABEX CORTEX and the FRM. E.P. and C.A. are employed by the Centre National de la Recherche Scientifique. We would like to thank Delphine Autran Clavagnier for help with data acquisition.

