## Supplementary Information for "Cognitive control of orofacial and vocal responses in the human frontal cortex"

**Supplementary Materials**

**Behavioral task perfoamce**

Three main performance measures were used from the learning and control tasks: 1) the proportion of correctly performed blocks, 2) the number of trials constituting the learning period (learning period length), and 3) the response time (RT) for each trial. With the proportion of correct blocks, we examined the effects of block type (learning/control), response type (manual/orofacial/vocal-verbal) and feedback type (vocal/verbal), and their possible interactions. A generalized linear mixed effect regression with binomial distribution was performed via the *glmer* function from the *lme* package (https://cran.r-project.org/web/packages/lme4/) in R statistical software. The statistical significance of the effects was assessed with Type II Wald tests implemented via the *Anova* function from the *car* package (https://cran.r-project.org/web/packages/car). For the trial RT data, we examined the effects of trial type (learning/post-learning/control), response type (manual/vocal/orofacial), and feedback type (vocal/verbal), and their possible interactions. Only behavioral data from the correctly performed learning and control task blocks were analyzed. A linear mixed effect regression with a normal distribution was performed via the *glmer* function. The statistical significance of the effects was assessed with Type II Wald tests implemented via the *Anova* function from the *car* package.

Linear mixed-model regression analyses revealed no significant effects of response type (*χ^2^*=3.22, *df*=3, *p*=0.36; Fig S1A) and feedback type (*χ^2^*=0.05, *df*=1, *p*=0.83, Fig S1A) on completion rates. This reflected that subjects performed equally well on the control and learning tasks when different responses were involved and when different feedbacks were provided. As expected, we observed a significant effect of task type (*χ^2^*=82.2, *df*=1, *p*<2x10^-16^, Fig S1A): the completion rates were higher in control versus learning blocks. As a further indication that response type had no influence on conditional-associative learning performance, we found that the mean learning period (number of trials taken to acquire the conditional associations in correctly performed blocks) did not differ across response types (*χ^2^*=3.05, *df*=2, *p*=0.218, Fig S1B).

In terms of response selection RTs, regression analyses revealed no significant effect of the type of feedback provided (*χ^2^*=2.02, *df*=1, *p*=0.16, Fig S1C). There was a significant main effect of response type (*χ^2^*=2.8x10^3^, *df*=3, *p*<2.2x10^-16^): subjects were fastest with manual responses, followed by orofacial, and non-speech vocal/verbal responses. Note that RTs in the visuo-vocal and visuo-verbal conditional learning association tasks were not significantly different (*p=1.00*). As expected, RTs were faster in control than learning trials, and in learning than post-learning trials (*χ^2^*=1.5x10^3^, *df*=2, *p*<2.2x10^-16^). Lastly, there was a significant interaction (*χ^2^*=63.4, *df*=6, *p*<8.9x10^-12^) between the response modality and trial type on the RTs, indicating that the effect of trial type (learning, post-learning, control trials) on RTs differed across the response types (manual, orofacial, nonspeech vocal, verbal). Post-hoc comparisons confirmed that, while the main effect of trial type was present in each response modality (all *p*<0.05) (i.e. control RTs are faster than learning RTs, and learning RTs are faster than post-learning RTs), the magnitude of differences between the trial types differed with response types: the difference between post-learning and control trials was the smallest for nonspeech vocal and verbal responses (*t*=12.7, 12.3, respectively, *p*<0.0001), followed by orofacial (*t*=21.9, *p*<0.0001) and manual responses (*t*=24.3, *p*<0.0001). This indicated that cognitive response selection based on learnt conditional associations appeared to be more efficient (as indicated by smaller increase in RTs from control) for vocal-verbal, followed by orofacial and manual responses. Similarly, the RTs differences between learning and control trials appeared to be the smallest for nonspeech vocal and verbal responses (*t*=10.6, 10.7, respectively, *p*<0.0001), orofacial responses (*t*=12.3, *p*<0.0001), and manual (*t*=21.5, *p*<0.0001) responses. This suggested that cognitive response selection during conditional associative learning appeared to be also the most efficient (as indicated by smaller increase in RTs from control) for vocal-verbal, followed by orofacial and manual responses.

**Supplementary Tables**

|  | **Learning minus Control** | | | | **Post-Learning minus Control** | | | |
| --- | --- | --- | --- | --- | --- | --- | --- | --- |
|  | *x* | *y* | *z* | *t* | *x* | *y* | *z* | *t* |
| **Manual** |  |  |  |  |  |  |  |  |
| L. PMd | -26 | -4 | 54 | 8.79 | -56 | 8 | -2 | 5.64 |
| L. BA44 (Dorsal) | -54 | 8 | 18 | 8.36 | - | - | - | - |
| L. BA44 (Ventral) | - | - | - | - | - | - | - | - |
| **Orofacial** |  |  |  |  |  |  |  |  |
| L. PMd | -26 | -2 | 50 | 4.19 | -30 | -8 | 52 | 6.48 |
| L. BA44 (Dorsal) | -54 | 14 | 20 | 6.58 | - | - | - | - |
| L. BA44 (Ventral) | -52 | 12 | 10 | 11.0 | -54 | 10 | -4 | 5.85 |
| **Vocal/Verbal** |  |  |  |  |  |  |  |  |
| L. PMd | - | - | - | - | - | - | - | - |
| L. BA44 (Dorsal) | -50 | 14 | 24 | 6.26 | - | - | - | - |
| L. BA44 (Ventral) | -52 | 18 | 6 | 4.49 | -56 | 8 | -2 | 5.64 |
| L. BA 45 | -48 | 32 | -2 | 4.69 | - | - | - | - |

**Table S1**– Group level analysis. Foci of increased activity in the posterior lateral frontal cortex observed during manual, orofacial, and vocal/verbal conditional selection in the learning and post-learning periods compared to their respective control conditions, i.e. during manual, orofacial, and vocal/verbal control selection. The x, y, z, coordinates are in MNI stereotaxic space.T-statistics are significant at *p_corrected_*<0.05.

|  | **Hemispheres with a pcgs** | | | | **Hemispheres without a pcgs** | | | |
| --- | --- | --- | --- | --- | --- | --- | --- | --- |
|  | *x* | *y* | *z* | *t* | *x* | *y* | *z* | *t* |
| **Manual** |  |  |  |  |  |  |  |  |
| MCC | \|4\| | 14 | 52 | 12.93 | \|2\| | 14 | 50 | 13.95 |
| MCC | \|10\| | 18 | 44 | 8.23 | \|4\| | 22 | 38 | 10.35 |
| Pre-SMA | - | - | - | - | - | - | - | - |
| **Orofacial** |  |  |  |  |  |  |  |  |
| MCC | \|2\| | 14 | 50 | 6.92 | \|6\| | 14 | 50 | 5.81 |
| MCC | \|2\| | 24 | 38 | 8.36 | \|6\| | 24 | 42 | 5.56 |
| Pre-SMA | - | - | - | - | - | - | - | - |
| **Verbal** |  |  |  |  |  |  |  |  |
| MCC | \|2\| | 14 | 54 | 5.94 | \|2\| | 18 | 50 | 4.17 |
| MCC | \|2\| | 26 | 46 | 6.22 | \|4\| | 26 | 40 | 4.65 |
| Pre-SMA | \|4\| | 8 | 68 | 4.52 | \|4\| | 8 | 72 | 4.16 |
| **Vocal** |  |  |  |  |  |  |  |  |
| MCC | \|4\| | 16 | 50 | 6.60 | \|2\| | 16 | 50 | 3.62 |
| MCC | \|6\| | 26 | 38 | 3.72 | \|4\| | 24 | 44 | 3.32 |
| Pre-SMA | - | - | - | - | - | - | - | - |

**Table S2**–Increased activity in the medial frontal cortex observed during manual, orofacial, vocal, and verbal conditional selection in learning period compared to that observed during manual, orofacial, vocal, and verbal control selection in hemispheres with and without a pcgs. X, Y, and Z coordinates correspond to the coordinates of the increased activities in the MNI stereotaxic space. T-statistics are significant at *p_corrected_*<0.05.

|  | **Verbal FB during learning minus control** | | | | **Vocal FB during learning**  **minus control** | | | |
| --- | --- | --- | --- | --- | --- | --- | --- | --- |
|  | *x* | *y* | *z* | *t* | *x* | *y* | *z* | *t* |
| **Manual** |  |  |  |  |  |  |  |  |
| *Left hemisphere* |  |  |  |  |  |  |  |  |
| PMd | -32 | -4 | 66 | 5.18 | -32 | -4 | 64 | 4.31 |
| MCC | 0 | 14 | 50 | 9.93 | 0 | 14 | 50 | 9.31 |
| MCC | -8 | 22 | 46 | 7.33 | -8 | 22 | 42 | 8.35 |
| BA 44 (Ventral) | -50 | 12 | -4 | 5.20 | -58 | 10 | 10 | 3.58 |
| *Right hemisphere* |  |  |  |  |  |  |  |  |
| MCC | - | - | - | - | 10 | 20 | 48 | 5.84 |
| **Orofacial** |  |  |  |  |  |  |  |  |
| *Left hemisphere* |  |  |  |  |  |  |  |  |
| MCC | -6 | 20 | 44 | 5.30 | -8 | 18 | 48 | 5.13 |
| Pre-SMA | -4 | 4 | 66 | 5.12 | - | - | - | - |
| BA 44 (Ventral) | 48 | 16 | 8 | 4.53 | -56 | 14 | 4 | 3.34 |
| *Right hemisphere* |  |  |  |  |  |  |  |  |
| MCC | 4 | 18 | 48 | 4.77 | 6 | 22 | 48 | 4.80 |
| **Verbal** |  |  |  |  |  |  |  |  |
| *Left hemisphere* |  |  |  |  |  |  |  |  |
| MCC | -6 | 18 | 44 | 5.18 | -4 | 16 | 50 | 5.40 |
| Pre-SMA | -6 | 10 | 64 | 4.87 | - | - | - | - |
| BA 44 (Ventral) | -50 | 12 | 0 | 3.35 | -52 | 12 | 2 | 4.04 |
| *Right hemisphere* |  |  |  |  |  |  |  |  |
| MCC | 8 | 20 | 48 | 4.50 | 8 | 20 | 46 | 4.15 |
| **Vocal** |  |  |  |  |  |  |  |  |
| Left hemisphere |  |  |  |  |  |  |  |  |
| MCC | -6 | 18 | 44 | 5.18 | -4 | 16 | 50 | 5.40 |
| Pre-SMA | -6 | 10 | 64 | 4.87 | - | - | - | - |
| BA 44 (Ventral) | -50 | 12 | 0 | 3.35 | -52 | 12 | 2 | 4.04 |
| Right hemisphere |  |  |  |  |  |  |  |  |
| MCC | 8 | 20 | 48 | 4.50 | 8 | 20 | 46 | 4.15 |

**Table S3**– Group level analysis. Increased activity in the posterior lateral frontal cortex observed during manual, orofacial, and vocal/verbal conditional selection in learning and post-learning periods compared to that observed during manual, orofacial, and vocal/verbal control selection. X, Y, and Z coordinates correspond to the coordinates of the increased activities in the MNI stereotaxic space. T-statistics are significant at *p_corrected_*<0.05.

|  | **Hemispheres with a pcgs** | | | | **Hemispheres without a pcgs** | | | |
| --- | --- | --- | --- | --- | --- | --- | --- | --- |
|  | *x* | *y* | *z* | *t* | *x* | *y* | *z* | *t* |
| **Verbal FB / Manual effector** | | | | | | | | |
| MCC | \|10\| | 20 | 48 | 7.22 | \|6\| | 18 | 48 | 8.46 |
| Pre-SMA | - | - | - | - | - | - | - | - |
| **Vocal FB / Manual effector** | | | | | | | | |
| MCC | \|2\| | 16 | 50 | 7.20 | \|8\| | 22 | 46 | 7.48 |
| Pre-SMA | - | - | - | - | - | - | - | - |
| **Verbal FB / Orofacial effector** | | | | | | | | |
| MCC | \|6\| | 18 | 46 | 7.93 | \|2\| | 18 | 48 | 4.22 |
|  | \|6\| | 30 | 34 | 5.91 |  |  |  |  |
| Pre-SMA | \|6\| | 8 | 62 | 3.96 | \|4\| | 4 | 66 | 3.27 |
| **Vocal FB / Orofacial effector** | | | | | | | | |
| MCC | \|6\| | 20 | 46 | 4.79 | \|2\| | 16 | 50 | 4.18 |
| Pre-SMA | - | - | - | - | - | - | - | - |
| **Verbal FB / Verbal effector** | | | | | | | | |
| MCC | \|6\| | 16 | 46 | 7.33 | \|8\| | 20 | 46 | 4.10 |
| Pre-SMA | \|4\| | 6 | 64 | 4.05 | \|4\| | 10 | 64 | 4.48 |
| **Vocal FB / Vocal effector** | | | | | | | | |
| MCC | \|6\| | 10 | 56 | 7.00 |  |  |  |  |
| MCC | \|6\| | 16 | 48 | 6.02 | \|6\| | 18 | 46 | 4.24 |
| Pre-SMA | - | - | - | - | - | - | - | - |

**Table S4**–Increased activity in the medial frontal cortex observed during vocal and verbal feedback analysis in visuo-manual, visuo-orofacial, visuo-vocal, and visuo-verbal conditional learning period compared to that observed during vocal and verbal feedback obtained in manual, orofacial, vocal, and verbal control trials in hemispheres with and without a pcgs. X, Y, and Z coordinates correspond to the coordinates of the increased activities in the MNI stereotaxic space. T-statistics are significant at *p_corrected_*<0.05.

**Supplementary Figure Legends**

**Figure S1. Behavioral data. A.** Proportion of blocks successfully completed in the conditional associative learning and control tasks displayed by response types (MAN-manual, ORO-orofacial, VE-verbal and VO-vocal) and feedback types (verbal – light blue bars, vocal – dark blue bars). **B.** Mean number of trials in the learning period in the manual (red), orofacial (orange) and vocal-verbal (yellow) response task versions. **C.** Mean reaction times in control (green bars), learning (orange bars) and post-learning (red bars) trials displayed by response types (MAN-manual, ORO-orofacial, VE-verbal and VO-vocal).
