## Supplementary figures and images for "Cognitive control of orofacial and vocal responses in the human frontal cortex"

### Supplementary Figures

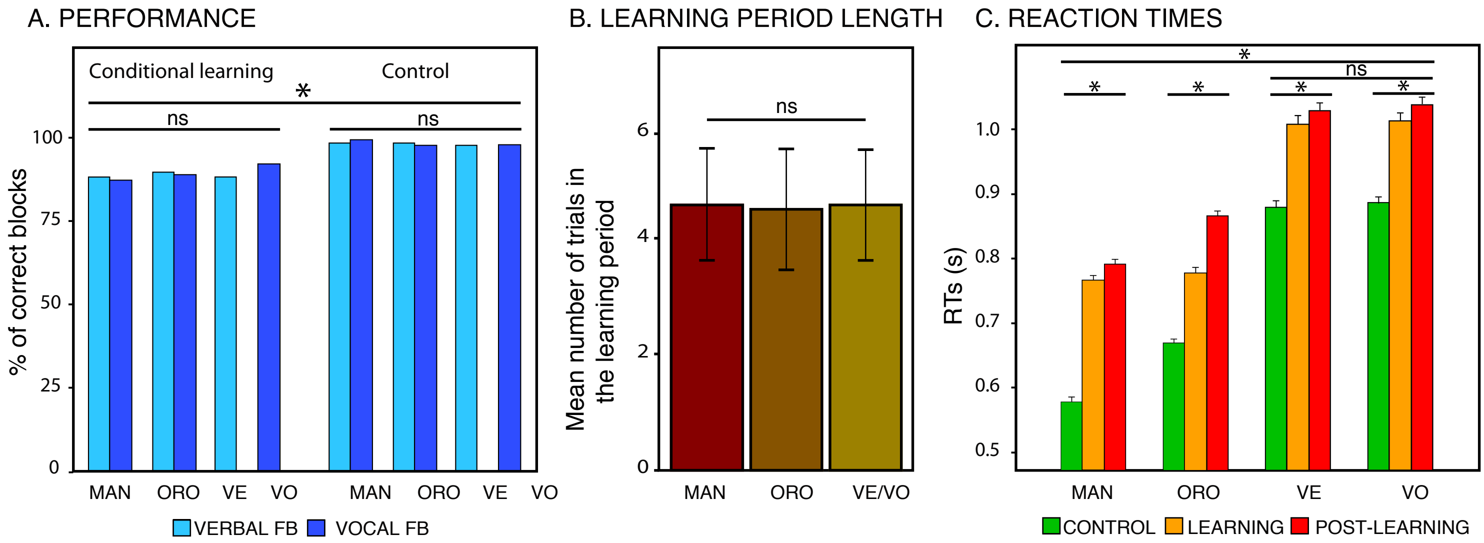
